# Deep sequencing of small RNA facilitates tissue and sex specific microRNA discovery in zebrafish

**DOI:** 10.1101/019760

**Authors:** Candida Vaz, Choon Wei Wee, Gek Ping Serene Lee, Philip W Ingham, Vivek Tanavde, Sinnakaruppan Mathavan

## Abstract

Role of microRNAs in gene regulation has been well established. Though the number of genes appear to be equal between human and zebrafish, miRNAs detected in zebrafish (∼247) is significantly low compared to human (∼2000; miRBase Release 19). It appears that most of the miRNAs in zebrafish are yet to be discovered. Using next generation sequencing technology, we sequenced small RNAs from brain, gut, liver, ovary, testis, eye, heart and embryo of zebrafish. In few tissues (brain, gut, liver) sequencing was done sex specifically. About 16-62% of the sequenced reads mapped to known miRNAs of zebrafish, with the exceptions of ovary (5.7%) and testis (7.8%). We used miRDeep2, the miRNA predication tool, to discover the novel miRNAs using the un-annotated reads that ranged from 7.6 to 23.0%, with exceptions of ovary (51.4%) and testis (55.2%) that had the largest pool of un-annotated reads. The prediction tool identified a total of 459 novel pre-miRNAs. Comparison of miRNA expression data of the tissues showed the presence of tissue and sex specific miRNAs that could serve as biomarkers. The brain and liver had highest number of tissue specific (36) and sex specific (34) miRNAs, respectively. Taken together, we have made a comprehensive approach to identify tissue and sex specific miRNAs from zebrafish. Further, we have discovered 459 novel pre-miRNAs (∼30% homology to human miRNA) as additional genomic resource for zebrafish. This resource can facilitate further investigations to understand miRNA-mRNA gene regulatory network in zebrafish which will have implications to understand human miRNA function.

## Introduction

The zebrafish has acquired scientific importance over the years and has become a powerful tool for unravelling the role of human genes. There are several reasons that make zebrafish as one of the most sought out organisms for biomedical research. High fecundity, fast growth rate, short generation time, ease of maintenance and an abundant supply of research material are some of the reasons for considering this model. Further, the availability of a number of genomic tools and the transparent embryos facilitate the non invasive strategy to observe experimental effects (Wixon 2000). Most importantly, comparison of human reference genome shows that at least 70% of the genes have one zebrafish orthologue (Howe et al. 2013). Of the human genes identified with morbidity descriptions in Mendelian Inheritance in Man (OMIM) database, about 82% can be related to at least one zebrafish orthologue (Howe et al. 2013). This high genomic similarity between zebrafish and human has resulted in important discoveries in the areas of human diseases, drug screens and therapeutic measures (Dodd et al. 2000); (Kari et al. 2007); (Goldsmith and Jobin 2012).

MicroRNAs (miRNAs) are a major class of non-coding RNA molecules that have steadily gained impetus over the past decade. They regulate several genes post-transcriptionally and serve as an added layer of control, being termed as “Micromanagers of gene expression” (Bartel and Chen 2004). The expression patterns of miRNAs represent the physiological state of the cell and thus have an essential prognostic capacity (Pritchard et al. 2012). Previous studies have revealed that comprehensive expression profiling of miRNA is useful in determining their specific expression patterns in cells (Lagos-Quintana et al. 2002), disease conditions such as cancer (Di Leva and Croce 2013), and during cell differentiation (Miska 2005); (Shi and Jin 2009).

There are several experimental methods for profiling miRNAs such as northern blotting (Sempere et al. 2004), RT-PCR (Chen et al. 2005a), microarrays (Sun et al. 2004) and RAKE assay (Berezikov et al. 2006). Each of these methods has their own limitations and advantages. Some of these methods are either cumbersome for scaling up and some are not sensitive enough to detect the low expressing miRNAs. The short length of these miRNAs also creates technical difficulties in the detection of mature miRNAs; further the above techniques are not suited to discover new miRNAs from the system.

With the advent of Next Generation sequencing (NGS) technology, there has been an increase in the efficiency of miRNA discovery. It not only overcomes the limitations of other methods but also provides an absolute and quantitative expression measurement (Hoen et al. 2008); (Creighton et al. 2009). NGS technology facilitates profiling of known miRNA and discovery of new transcripts. Owing to its high sensitivity, this technology can identify low abundant miRNA transcripts, which may not be detected by other methods.

In zebrafish, miRNA expression profiling and discovery was initially carried out through small RNA cloning and microarray analysis. MicroRNA expression was detected through large scale sequencing of sRNA libraries prepared from different developmental stages of zebrafish and two adult cell lines (Chen et al. 2005b) and this study identified about 154 mature miRNAs. Further this study revealed that the early zygotic stage (0h) stage is nearly devoid of miRNAs. The expression of miR-430 family peaks at the blastula stage (4h) and dominates the miRNA profile up to the pharyngula stage (24h) and then decreases. Role of miR-430 in maternal RNA clearance during maternal to zygotic transition has been well documented (Giraldez et al. 2006). Kloosterman and co-workers identified 139 known and 66 new miRNA from 5-day old zebrafish larvae and adult zebrafish brain; using in situ hybridization and northern blotting they identified developmental stage specific and tissue specific expression patterns for some of these miRNAs (Kloosterman et al. 2006).

Using pyrosequencing technology to discover miRNAs, Soares et al retrieved 90% of the known miRNAs and predicted 25 novel miRNAs from different developmental stages and fully developed organs of zebrafish (Soares et al. 2009).

One of the recent publications used NGS technology to determine the temporal expression patterns for miRNAs and piRNAs during early embryonic development of zebrafish (Wei et al. 2012). In this study, they identified a number of known miRNA, 8 novel miRNAs and a diverse set of piRNAs (Wei et al. 2012). Based on all the above mentioned studies, a total number of 247 mature and 344 pre-miRNAs were identified and submitted as genomic resource for zebrafish (miRBase Release 19). However, no attempt has been made to identify tissue specific or sex specific miRNAs from the known miRNA resource. MicroRNA database (miRBase) (Griffiths-Jones 2004); (Griffiths-Jones et al. 2006); (Griffiths-Jones et al. 2008) has a total of ∼2000 mature miRNAs for human and only about 247 miRNAs for zebrafish. It is known that number of coding genes in human and zebrafish are almost same. However, zebrafish miRBase has very low number of miRNA compared to human. Thus it appears that most of the miRNA in zebrafish are yet to be discovered.

Using NGS technology we undertook a systematic and comprehensive approach to determine the tissue/sex specific expression patterns of known miRNAs and to discover novel miRNAs from different tissues of zebrafish such as the brain, gut, liver, ovary, testis, eye, heart and embryo. For tissues such as brain, gut and liver, smallRNA from both the male and female counterparts were sequenced separately. The aim of our study was to discover new miRNAs from different tissues and to identify tissue specific and sex specific expression of known and novel miRNAs. Our results are highly reliable because each sample was sequenced using 3 biological replicates and generated approximately 20M reads/per library. Thus our small RNA deep sequencing data had a high sequencing depth (4-10 times more than the required depth) that is deep enough for expression profiling of even the lowly expressed miRNAs and for discovery applications.

The expression pattern of miRNAs specific to each tissue and gender can serve as a useful resource for further research and for understanding the miRNA-mRNA regulatory mechanisms in zebrafish. We have also discovered about 459 putative novel pre-miRNAs, of which about 30% had human homologs and rest are zebrafish specific. (Griffiths-Jones 2004); (Griffiths-Jones et al. 2006); (Griffiths-Jones et al. 2008); (Kozomara and Griffiths-Jones 2011); (Kozomara and Griffiths-Jones 2014). This study adds a significant genomic resource to the zebrafish miRNA landscape and has potential to provide better understanding of miRNA based gene regulation in zebrafish.

## Results

### Micro RNA frequency distribution in the sRNA datasets

Zebrafish small RNA deep sequencing data from different tissues was generated using Illumina HiSeq 2000 platform. Each library comprised of three biological replicates and approximately 20 million reads/library. The details of the number of reads for each library are shown in **Table 1**. Datasets from each miRNA library (three biological replicates) generated in this work were matched to different annotated databases to classify the sRNA sequenced reads into different RNA categories (details in the “Methods” section) to get an overview of the frequency of different classes of RNA present in the samples as well as to obtain the unannotated pool of sequenced reads for novel miRNA prediction. The flowchart depicting the entire analysis pipeline is shown in **Figure 1**. In general, the sequenced reads small RNA that mapped to the known miRNA database was abundant in female tissues compared to male tissues (female: brain 62.2%; gut 37.0%; liver 35.8%) (male: brain 38.2%; gut 33.1%; liver 20.8%). Expression for eye and heart was tested only in male tissues and it amounted to 33.5% and 34.5%, respectively. The sequenced reads mapping to known miRNA was low in embryo (16.3%), ovary (5.7%) and testis (7.8%) and these samples had large number of unannotated reads.

**Table 1.** Details of the zebrafish sRNA deep sequencing datasets. Each tissue library comprised of three biological replicates. One of the replicate of the heart did not cluster well with the other replicates and was not considered for further analysis. The table shows the number of processed (adaptor trimmed, >=18nt) reads for each biological replicate. Around 92-98% of reads mapped to the zebrafish genome.

| Library | Replicates | Total number of reads | Total number of reads (in millions) | Percentage of reads mapped |
| --- | --- | --- | --- | --- |
| Embryo | Embryo1 | 7565225 | 7.6 | 94.4 |
|  | Embryo2 | 10202969 | 10.2 | 94.8 |
|  | Embryo3 | 32247131 | 32.2 | 94.8 |
| Male Brain | Male Brain1 | 5101418 | 5.1 | 97.8 |
|  | Male Brain2 | 6533205 | 6.5 | 97.3 |
|  | Male Brain3 | 10518347 | 10.5 | 97.0 |
| Female Brain | Female Brain1 | 2338429 | 2.3 | 96.7 |
|  | Female Brain2 | 3278013 | 3.3 | 97.5 |
|  | Female Brain3 | 12648618 | 12.6 | 97.8 |
| Male Gut | Male Gut1 | 16232210 | 16.2 | 97.1 |
|  | Male Gut2 | 8864108 | 8.9 | 97.7 |
|  | Male Gut3 | 3117877 | 3.1 | 95.8 |
| Female Gut | Female Gut1 | 13939922 | 13.9 | 97.4 |
|  | Female Gut2 | 8260495 | 8.3 | 97.8 |
|  | Female Gut3 | 7493835 | 7.5 | 97.9 |
| Male Liver | Male Liver1 | 18404267 | 18.4 | 97.8 |
|  | Male Liver2 | 9369428 | 9.4 | 97.4 |
|  | Male Liver3 | 4639550 | 4.6 | 97.1 |
| Female Liver | Female Liver1 | 5060891 | 5.1 | 97.0 |
|  | Female Liver2 | 8162716 | 8.2 | 97.2 |
|  | Female Liver3 | 1646108 | 1.6 | 96.4 |
| Ovary | Ovary1 | 8824267 | 8.8 | 95.8 |
|  | Ovary2 | 14310754 | 14.3 | 95.9 |
|  | Ovary3 | 25898221 | 25.9 | 95.9 |
| Testis | Testis1 | 17427858 | 17.4 | 92.5 |
|  | Testis2 | 20235869 | 20.2 | 92.2 |
|  | Testis3 | 14203535 | 14.2 | 92.5 |
| Eye | Eye1 | 9398588 | 9.4 | 96.0 |
|  | Eye2 | 5722393 | 5.7 | 95.5 |
|  | Eye3 | 2854316 | 2.9 | 95.9 |
| Heart | Heart1 | 38070247 | 38.1 | 95.2 |
|  | Heart2 | 34828065 | 34.8 | 96.2 |

**Figure 1.**
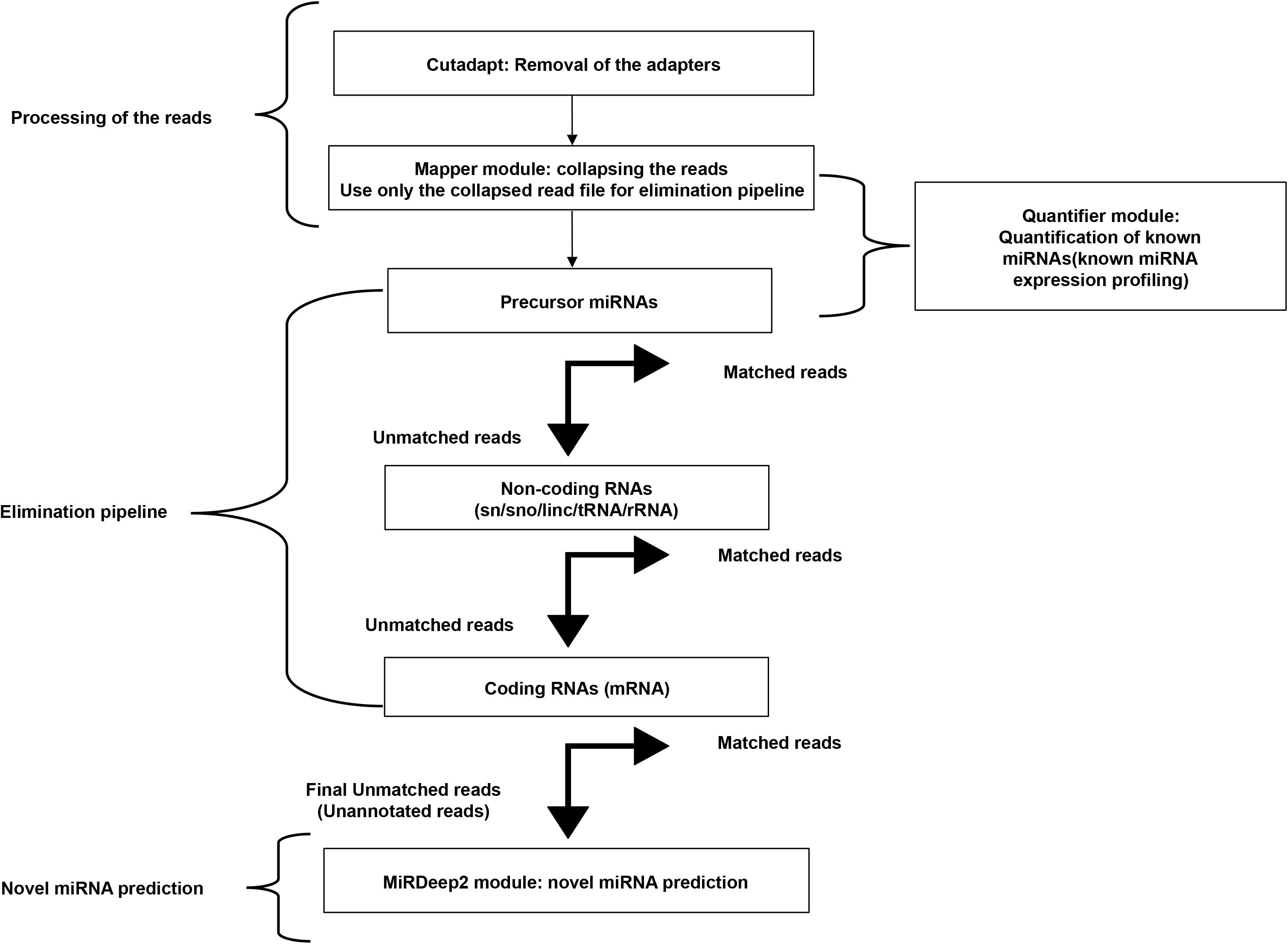
Flowchart depicting the workflow for known miRNA profiling and novel miRNA prediction. The workflow comprises of four parts namely: Processing of the reads, Quantification of known miRNAs, the Elimination pipeline and the Novel miRNA prediction pipeline. For the elimination pipeline, the reads were matched with the known annotated genomic sequences. Alignment with maximum of two mismatches was considered as matched reads. All the matched reads were removed before the next round of elimination. The quantification of known miRNAs and prediction of novel miRNAs was done using miRDeep2 modules.

Small RNA seq data contained significant representation of tRNAs in most of the tissues, with the maximum amount in the gut (male 45.8%; female: 45.1%) and liver (male 53.2%; female 42.6%) **(Figure 2)**. The mRNAs (small size/degraded/fragmented coding RNA) were more abundant in smallRNA seq data of the embryo and reproductive tissues (embryo 23.1%; ovary 20.1% and testis 16.0%) than in others. rRNAs and other non-coding RNAs were the minimal RNA species in all the datasets ranging from 0.3-4.5% and 1-5.4% respectively **(Figure 2)**. The percentage of un-annotated mapped reads ranged from 7.6 to 23.0% with exception of the ovary (51.4%) and testis (55.2%). The sequences that did not map to the genome ranged from 2.3 to 7.6% **(Figure 2).**

**Figure 2.**
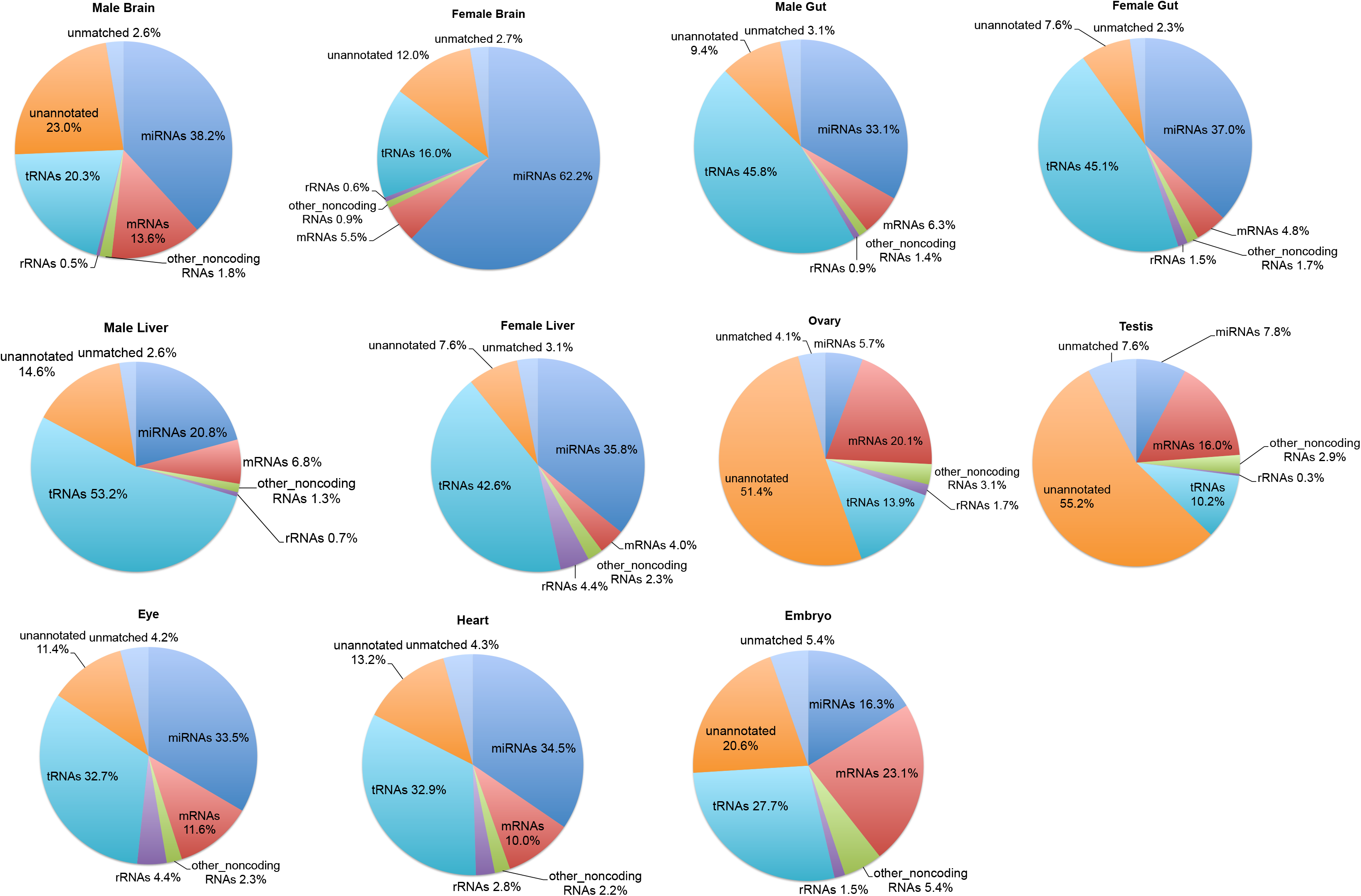
Pie charts depicting the frequency of different classes of RNA species present in the sRNA datasets. The pie charts represent an overview of the distribution of small RNAs in the different tissue libraries. For the male brain and female brain, the miRNAs constitute the most abundant sRNA, for the male gut, female gut, male liver and female liver, the tRNAs constitute the most abundant sRNA, and for the eye and heart, the miRNAs and tRNAs equally constitute the most abundant sRNAs. The ovary, testis and embryo had comparatively lesser amount of miRNAs. The ovary and testis had nearly half of their sRNA reads unannotated.

### Expression pattern of known miRNAs

Out of the 247 known mature miRNAs, 86-96% were detected by the quantifier module of miRdeep2 (Friedlander et al. 2012) as they were found to be expressed in one or the other tissue sample (See “Methods”) (**Supplemental File S1).** The raw expression data/raw read counts of the mature miRNAs of replicates of each sample were pooled and binned. The known mature miRNAs showed a wide range of expression values spanning up to 7 levels of magnitude (**Level 1**: 1-10, **2**: 10-10^2^, **3**: 10^2^-10^3^, **4**: 10^3^-10^4^, **5**: 10^4^-10^5^, **6**: 10^5^-10^6^, **7**:10^6^-10^7^) (**Figure 3A**). The expression pattern of the known mature miRNAs of all tissue samples showed almost normal distribution with the peak at Level 4. Maximum miRNAs had expression values within Level 4 (10^3^-10^4^), followed by Level 3 (10^2^-10^3^), Level 5 (10^4^-10^5^) and Level 2 (10-10^2^) ranges, respectively. Few miRNAs occurred in the lowest level, Level 1 (1-10) and second highest level, Level 6 (10^5^-10^6^), with the minimum at the highest level, Level 7 (10^6^-10^7^) (**Figure 3A**).

**Figure 3.**
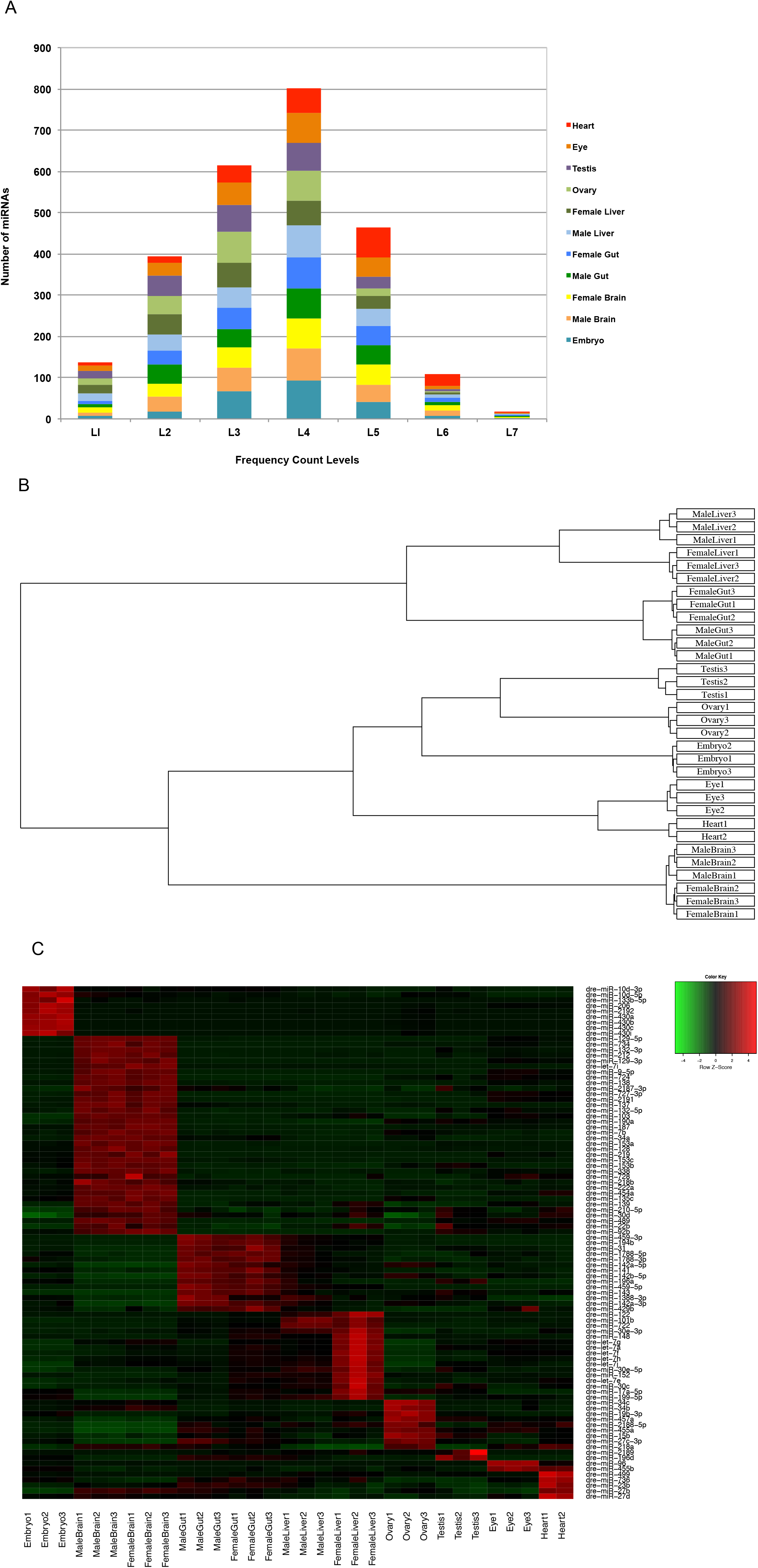
Tissue specific expression pattern of the known miRNAs. **A – Distribution of known miRNA expression levels with respect to number of miRNAs.** Numbers of reads are taken as miRNA expression levels and their values are represented in the form of ranges: Level 1: 1-10, Level 2: 10-10^2^, Level 3: 10^2^-10^3^, Level 4: 10^3^-10^4^, Level 5: 10^4^-10^5^, Level 6: 10^5^-10^6^, Level 7:10^6^-10^7^. **B – Hierarchical clustering plot showing the similarity and differences among the samples.** The known mature miRNA profiles of all the tissues were first subjected to TMM normalization and their clustering pattern was determined. There are two major groups seen: (i) The gut and the liver samples (ii) The brain, heart, eye, embryo, ovary and testis samples. The second major group is further divided into three subgroups: (a) The ovary and testis samples were closely placed, followed by the embryo (b) The eye and heart samples (c) The brain samples **C – Expression profile of the tissue specific known miRNAs.** Differential expression of tissue specific known mature miRNAs. The upregulated miRNAs are depicted in red colour whereas the down regulated miRNAs are depicted in green colour.

### Clustering of the sRNA datasets

The known mature miRNA profiles of all the tissues were first subjected to Trimmed Mean of M-values (TMM) normalization (**Supplemental File S1**) using the Bioconductor package edgeR (Robinson et al. 2010) as mentioned in the “Methods” section and their clustering pattern was determined using the hierarchical clustering plot (**Figure 3B**). The data sets were also subjected to PCA analysis and clustering pattern obtained is presented in (**Supplemental Figure S1**). The three biological replicates of all the tissue samples including embryo clustered closely indicating less experimental variation or noise among the replicates. However, one of the replicates of the heart sample did not cluster with the other replicates and hence was removed from all further analysis. The tissue samples having male and female counterparts like the brain, gut and liver showed proximity among their male and female counterparts, indicating that the variations caused due to sex was less compared to the variations caused by the tissue. The samples were grouped as described in the hierarchical clustering plot (**Figure 3B**) and the PCA plot (**Supplemental Figure S1).**

### Tissue specific known miRNAs

To determine the tissue specific known miRNAs, the TMM normalized expression profiles of the known mature miRNAs of all the tissues were first compared with those of the embryo samples; using GLM edgeR analysis (Robinson et al. 2010) (see “Methods”) to obtain the differentially expressed miRNAs (DE miRNAs). The number of differentially expressed known mature miRNAs of the tissues with respect to the embryo is presented in **Supplemental Table S1**. The details of the known DE mature miRNAs of all the samples, including the logFC, logCPM, P-value and FDR are presented in the **Supplemental File S2**.

The up-regulated DE known mature miRNA sets of all the tissues were compared with each other except with its male/female counterpart to obtain the up-regulated DE miRNAs only found or the most enriched in that particular tissue (see “Methods”), or in other words, tissue specific miRNAs. The union of the tissue specific miRNAs for the male and female counterparts of brain, gut and liver were taken as their respective tissue specific miRNAs.

To obtain embryo specific known miRNAs, the down-regulated DE known mature miRNA sets of all the tissues were compared to obtain the commonly down regulated miRNAs. These commonly down regulated miRNAs were called “embryo specific”.

The brain had the highest number of known tissue specific miRNAs (36), followed by the liver (16), gut (14), ovary (9), heart (5), testis (2), and eye (2). The embryo had 9 specific miRNAs **(Supplemental Table S2, Figure 3C, Supplementary File S3)**.

A number of tissue specific miRNAs reported by earlier studies involving in situ hybridisation and Northern Blots (Kloosterman et al. 2006) were also detected in the current analysis in the respective tissues. For example, miRNAs such as : dre-miR-135c, dre-miR-454a, dre-miR-724-5p, dre-miR-727-3p, dre-miR-728 and dre-miR-734 were found to be specific or highly expressed in brain, dre-miR-459-3p found to be specific to anterior part of the gut, dre-miR-122 found to be specific to liver, dre-miR-455 found to be specific to eye and dre-miR-499 found to be highly enriched in heart. The dre-miR-430 miRNA family was specific for embryo (Kloosterman et al. 2006).

### Sex specific known miRNAs

To determine the sex specific known miRNAs, the TMM normalized expression profiles of known mature miRNAs of the male and female counterparts of the brain, gut and liver were compared using the GLM edgeR analysis (Robinson et al. 2010) to obtain DE mature miRNAs. The DE mature miRNAs among the male and female counterparts were deemed as the sex specific miRNAs. The liver had the highest number of sex specific miRNAs (34), followed by the brain (9) and gut (7) **(Supplemental Table S3 and Figure 4A-C).** Details of the sex specific miRNAs including their logFC, logCPM, P-value, FDR are given in **Supplemental File S4**. The miRNA dre-miR-2190 was found to be commonly sex-specific for brain, gut and liver, enriched in the male counterparts. The miRNAs dre-miR-34c, dre-miR-148, dre-miR-430a, dre-miR-430b, dre-miR-430c, were found to be commonly sex specific for gut and liver, enriched in the female counterparts. On comparison of the ovary and testis samples, a total of 63 miRNAs were found to be differentially expressed among them **(Figure 4D).** Details of the DE mature miRNAs among ovary and testis are given in **Supplemental File S4**. The miRNAs dre-miR-34c, dre-miR-430a, dre-miR-430b, dre-miR-430c were also enriched in the ovary as compared to the testis as seen for the female counterparts of the gut and liver

**Figure 4.**
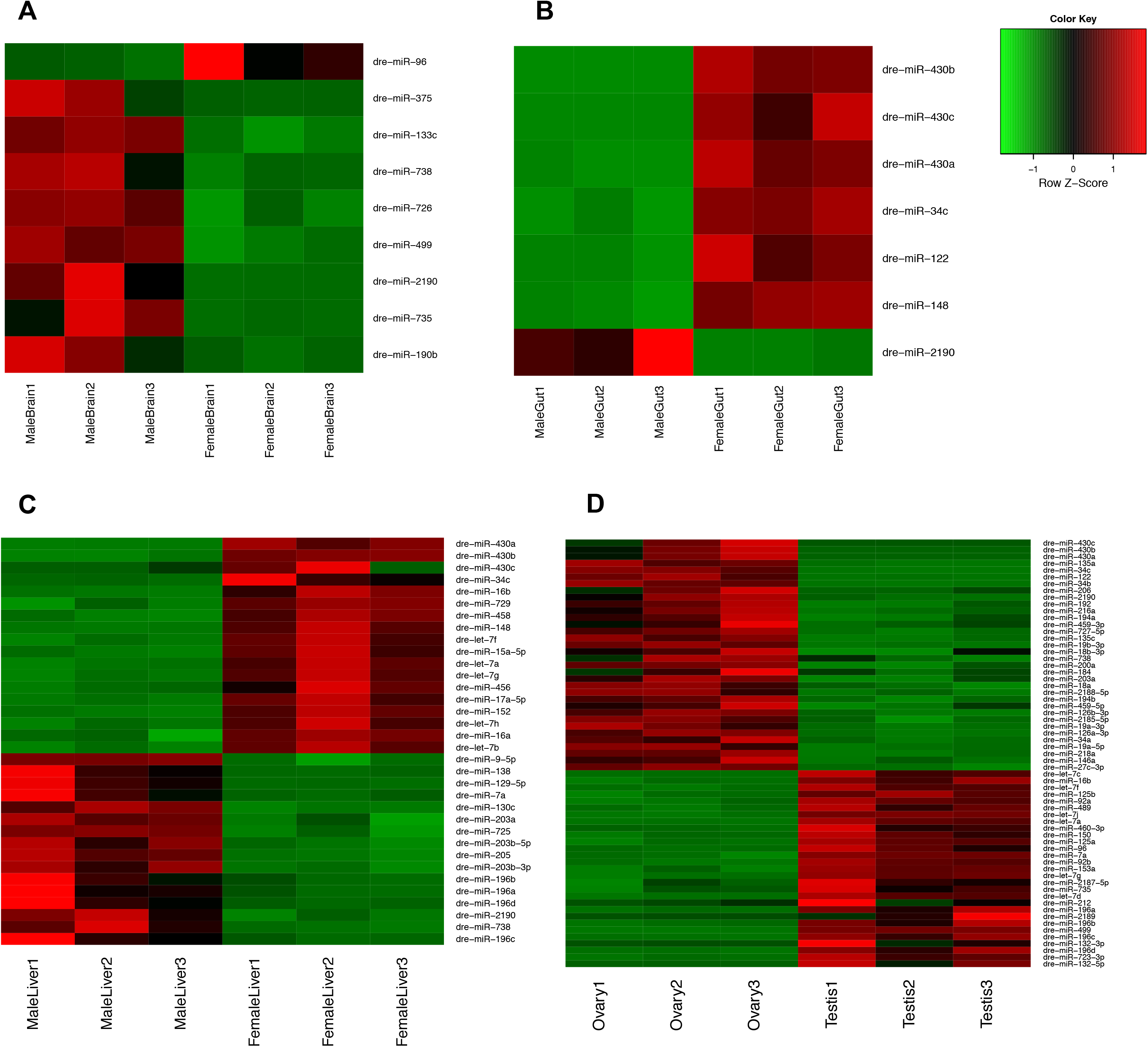
Expression profile of the sex specific known miRNAs. Differential expression of sex specific known mature miRNAs for (A) brain (B) gut (C) liver (D) ovary vs testis. The upregulated miRNAs are depicted in red colour whereas the down regulated miRNAs are depicted in green colour.

### Novel miRNA discovery

The reads that did not map to any of the known annotated data bases (un-annotated reads) were identified through the elimination pipeline and these un-annotated reads were used for novel miRNA prediction. The un-annotated sequences of the three replicates of a tissue were first combined together and then subjected to the miRDeep2.pl module (Friedlander et al. 2008); (Friedlander et al. 2012) for prediction of novel miRNAs. Since the three replicates were of different sequencing depths, it was essential to combine their un-annotated reads before running the miRDeep2 module to obtain consistent predictions for each tissue.

To test the sensitivity of the miRDeep2, each sample was run through the miRDeep2 before being sieved through the elimination pipeline. The number of known miRNAs detected by miRDeep2 (at its default cut-off) in comparison to the total number of miRNAs in the particular sample was indicative of the sensitivity of miRDeep2. The sensitivity of miRDeep2 ranged from 89% to 95% with the exception of one female liver sample, for which the sensitivity was 81% (**Supplemental Figure S2**).

The **Table 2** lists the number of novel pre-miRNAs predicted for each sample and the details of the predicted novel pre-miRNAs including the information regarding the miRdeep score, mature miRNA read count, miRNAs with same seed region, consensus mature miRNA sequence, consensus precursor miRNA sequence, precursor miRNA genomic coordinates, for each tissue sample is given in **Supplemental File S5**. The **Table 2** also shows the number of human homologs for the predicted novel miRNA. The structure of a predicted novel pre-miRNA along with the reads aligned to it is shown in **Supplemental Figure S3.**

**Table 2.**
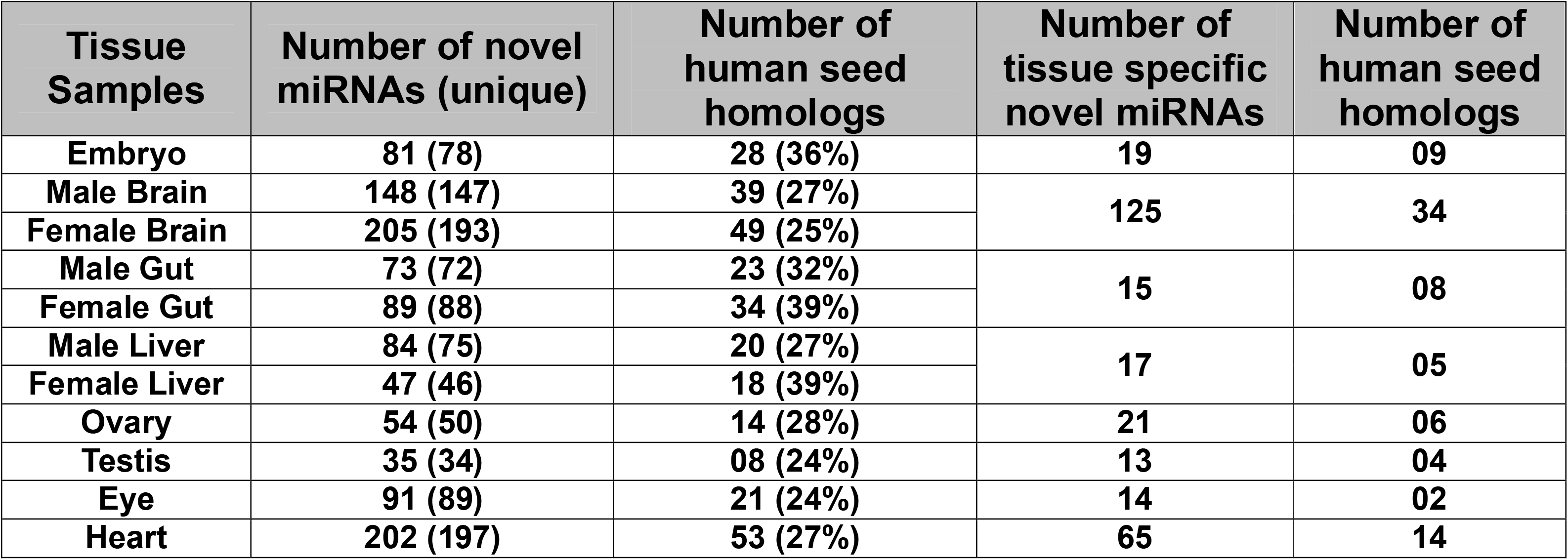
Details of the novel pre-miRNAs predicted from different tissues. The un-annotated sequences left after the elimination pipeline, were used for the novel miRNA prediction. For this purpose, the un-annotated sequence files of the replicates of a particular tissue were first combined and mapped to the zebrafish genome using mapper.pl module and then subjected to the miRDeep2.pl module to obtain the novel miRNAs. The table shows the total number of predicted novel pre-miRNAs as well as those that were specific for each tissue. The value in brackets is the unique number of novel pre-miRNAs predicted for each tissue. The miRDeep2.pl module also gives the seed homologs if present for each predicted novel miRNA. The table shows the number of predicted total and tissue specific novel miRNAs having human seed homologs. On an average 30% of the predicted novel pre-miRNAs had human seed homolgs.

| **Tissue Samples** | **Number of novel miRNAs (unique)** | **Number of human seed homologs** | **Number of tissue specific novel miRNAs** | **Number of human seed homologs** |
| --- | --- | --- | --- | --- |
| **Embryo** | **81 (78)** | **28 (36%)** | **19** | **09** |
| **Male Brain** | **148 (147)** | **39 (27%)** | **125** | **34** |
| **Female Brain** | **205 (193)** | **49 (25%)** |  |  |
| **Male Gut** | **73 (72)** | **23 (32%)** | **15** | **08** |
| **Female Gut** | **89 (88)** | **34 (39%)** |  |  |
| **Male Liver** | **84 (75)** | **20 (27%)** | **17** | **05** |
| **Female Liver** | **47 (46)** | **18 (39%)** |  |  |
| **Ovary** | **54 (50)** | **14 (28%)** | **21** | **06** |
| **Testis** | **35 (34)** | **08 (24%)** | **13** | **04** |
| **Eye** | **91 (89)** | **21 (24%)** | **14** | **02** |
| **Heart** | **202 (197)** | **53 (27%)** | **65** | **14** |

### Expression pattern of novel predicted miRNAs

The novel pre-miRNAs detected from all the tissues including the embryo sample were grouped together and collapsed to obtain a non redundant set of novel pre-miRNAs. The analyses predicted a final set of 459 unique novel pre-miRNAs and were given an identifier prefix: “gis-dre-mir”. **The Supplemental File S6** comprises of the precursor sequences of the 459 predicted novel miRNAs with their mature sequences in fasta format.

The predicted novel mature miRNAs showed a skewed distribution of expression unlike the known mature miRNAs with the peak at the lowest level Level 1: 1-10 and a decreasing trend toward the higher levels. There were almost no miRNAs at the highest levels Level 6: 10^5^-10^6^ and Level 7:10^6^-10^7^ (**Figure 5A**). This verifies the previous findings that the most abundant miRNAs are already known and the novel miRNAs tend to be rare and low in expression (Kloosterman et al. 2006).

**Figure 5.**
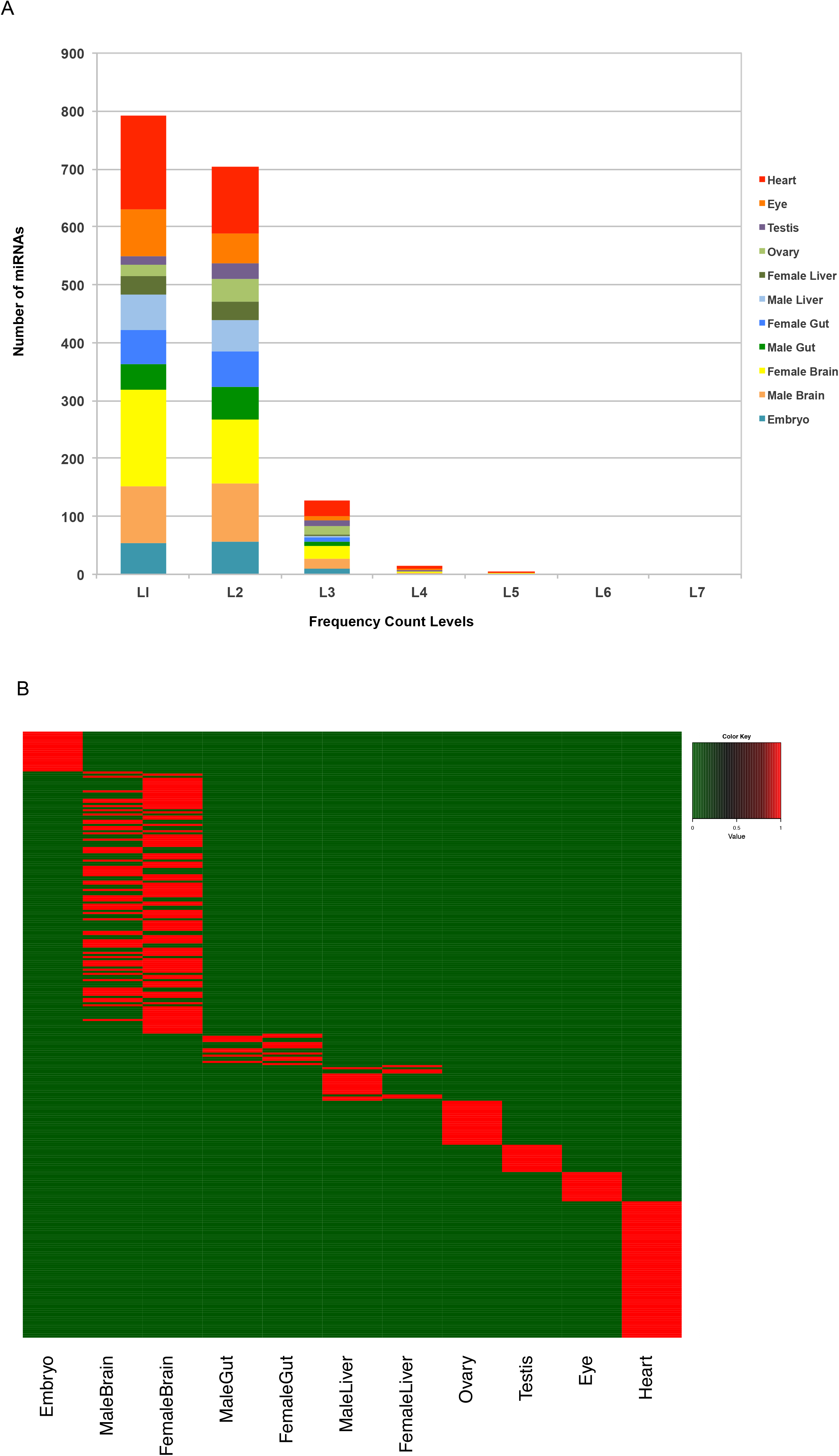
Tissue specific expression pattern of the novel miRNAs. **A – Distribution of novel miRNA expression levels with respect to number of miRNAs.** Numbers of reads are taken as miRNA expression levels and their values are represented in the form of ranges. Level 1: 1-10, Level 2: 10-10^2^, Level 3: 10^2^-10^3^, Level 4: 10^3^-10^4^, Level 5: 10^4^-10^5^, Level 6: 10^5^-10^6^, Level 7:10^6^-10^7^. **B – Expression profile of the tissue specific novel pre-miRNAs.** Differential expression of tissue specific novel pre-miRNAs. The presence of a novel pre-miRNA is depicted by red colour whereas the absence of it is depicted by green colour.

### Tissue specific novel pre-miRNAs

The novel pre-miRNAs were compared among all the samples including the embryo sample to obtain tissue-specific predicted novel pre-miRNAs as well as embryo specific predicted novel pre-miRNAs. **Supplemental Figure S4** gives the schematic representation of the method to obtain these tissue specific novel pre-miRNAs. The number of predicted tissue-specific novel pre-miRNAs for each sample is shown in **Table 2 and Figure 5B**. The brain showed the highest number of predicted tissue specific novel pre-miRNAs (125) followed by the heart (65). The details of these predicted tissue-specific novel pre-miRNAs are given in **Supplemental File S7**.

### qPCR validations

Some of the known and novel predicted miRNAs were validated through qPCR and their expression patterns were compared with the expression patterns obtained through miRNAseq. The expression pattern obtained by RNA seq and qPCR was compared for selected miRNAs as a validation strategy. A total of 8 known miRNAs and 13 novel miRNA were tested by qPCR, out of which 5/8 known and 12/13 novel miRNAs showed similar patterns with correlation values above 0.5. Mostly, those which have low tag counts did not show significant correlation between miRNA seq and qPCR (see “Methods”). All the 5 known miRNAs having correlation above 0.5 and the top 5 among the 12 novel miRNAs are shown in **Figure 6**.

**Figure 6.**
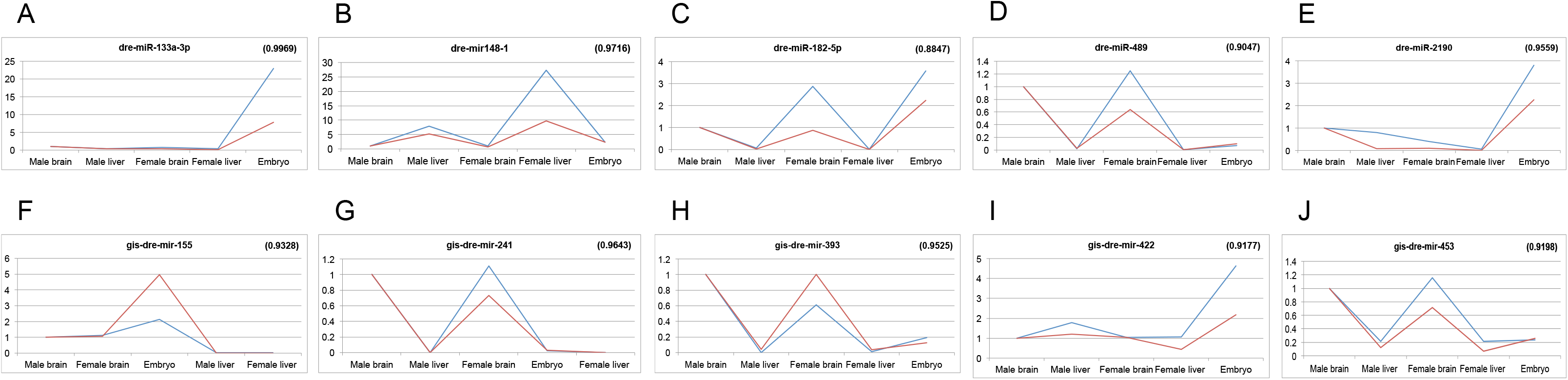
miRNA expression levels using qPCR and miRNA sequencing show a good correlation. (A-E) Known miRNAs (F-J) Novel miRNAs. Blue line denotes expression values based on qPCR and red line denotes expression values based on miRNA sequencing. A correlation comparison was done between the expression profiles from both platforms and the values are shown in brackets.

## Discussion and Conclusions

This study involves an exhaustive small RNA deep sequencing from several tissues and the embryo of zebrafish and systematic and comprehensive analysis of the data. The major focus was first to profile and identify tissue-specific and sex-specific known miRNAs and second to discover novel miRNAs.

Generally, a RNAseq tag density of 1–2M reads is good enough for miRNA expression profiling and, a tag density of 2–5M reads is sufficient for discovery applications. We have generated about 20M reads for each library in the current project, which is four to ten times more than the required range. Further we sequenced 3 biological replicates for most of the samples. Hence, our sRNA deep sequencing data had a high sequencing depth, deep enough for expression profiling and sensitive enough to discover lowly expressed novel miRNAs.

Around 92-98% of the reads mapped to the zebrafish genome (Zv9) indicating minimal amount of degradation in the samples. The known miRNAs (344 precursors, 247 mature) comprised a wide range of the mapped reads (16-62%), with the brain samples having the maximum amount (38-62%) and with the ovary and testis having the minimum amount (6-8%). The final un-annotated pool of sequences that serves as a source of novel miRNA prediction was 7.6 to 23.0% with exceptions of ovary (51.4%) and testis (55.2%) that had almost half of the sRNA sequences un-annotated. The presence of such a high amount of un-annotated sequences in ovary and testis suggests the presence of some unknown small non-coding RNAs such as piRNA that needs to be further analysed. It has been shown that piRNA are most abundant in reproductive tissues and early embryos (Cox et al. 2000); (Houwing et al. 2007).

The known mature miRNAs in each sample showed a wide range of expression values spanning 7 levels of magnitude. The expression values of the known miRNAs of all tissue samples showed an almost normal distribution.

To obtain tissue specific known miRNAs, the mature miRNA expression profile of each tissue sample was compared to the embryo to obtain the differentially expressed miRNAs. Since the embryo is undifferentiated and a precursor of the tissues it was taken as control. The entire zebrafish could also have been used a control, but since the embryo is a precursor of all tissues and more homogenous as compared to the whole fish, it was a better choice to use as a control for comparison to other tissues to obtain miRNAs that get de-regulated during differentiation. The miR-430 family was found to be very highly expressed in embryo with very low to none expression in adult tissues as seen in previous studies (Chen et al. 2005b). The miR-430 family has been shown to be involved in maternal RNA degradation during gastrulation (Giraldez et al. 2006). Since our samples contained gastrulation stage embryo as well, it is logical to expect abundant presence of miR-430 in the sequence data. This miRNA family was found to be commonly down regulated in all the adult tissues. Their specific expression in female liver and gut indicate that this miRNAs might be involved in functions in addition to maternal RNA degradation

The upregulated mature miRNAs of the tissues were then compared among each other to get the tissue specific known miRNAs. The brain showed the highest number of tissue specific known miRNAs (36) pointing out to the fact that these miRNAs may have significant roles in brain development and functioning. Among the tissue specific miRNAs detected, some of them were reported in previous studies (Kloosterman et al. 2006); (Soares et al. 2009).

The sex-specific miRNAs were obtained by comparing the mature miRNA expression profiles of the male and female counterparts of brain, gut and liver. The liver showed the highest number of sex-specific known miRNAs (34), revealing major differences in the functioning of male and female livers. The miRNA dre-miR-2190 was found to be commonly sex-specific for brain, gut and liver, enriched in the male counterparts. The miRNAs dre-miR-34c, dre-miR-148, dre-miR-430a, dre-miR-430b, dre-miR-430c, were found to be commonly sex specific for gut and liver, enriched in the female counterparts as well as ovary (on comparison with testis).

One of the most important applications of Next Generation Sequencing is the ability to detect novel miRNAs. The prediction of novel miRNAs was done by miRDeep2 software (Friedlander et al. 2008); (Friedlander et al. 2012) using the un-annotated pool of sequences, left after the removal of the annotated sequences through the elimination pipeline. The sieving out of the known and annotated sequences through the elimination pipeline cuts down on the number of false predictions and increases the specificity of novel miRNA prediction.

Before using miRDeep2 for novel miRNA prediction, we ran the software on the un-sieved and entire samples comprising of the annotated and unannotated reads. The number of known miRNA picked up by miRDeep2 in comparison to the total number of miRNAs in the samples at the similar cut-off used for novel miRNA prediction was used as an indicator of sensitivity. The sensitivity of miRDeep2 ranged from 89 to 95% with the exception of one female liver sample, for which the sensitivity was 81%.

A total of 459 novel pre-miRNAs were predicted in this study based on the sequence data. Majority of these novel miRNAs were less abundant and showed a skewed frequency distribution, with large number of miRNAs falling within the low levels of expression and very few within the high levels of expression. This is perhaps the reason they have not been discovered so far and highlights the merit of our really deep sequencing approach. These predicted novel miRNAs were also found to be having less seed conservation to human miRNAs (30% on an average) and thus indicating the most of the novel miRNAs are zebrafish specific. These findings were in accordance to an earlier study that also reported less abundance and less conservation of the novel miRNAs (Kloosterman et al. 2006).

The sequences of the predicted novel pre-miRNAs were compared with each other to obtain the ones specific to each tissue. Here again, the brain showed maximum number (125) tissue specific novel pre-miRNAs.

It would be further interesting to correlate these tissue and sex specific miRNAs to their targets to understand the gene regulatory network and pathways regulating zebrafish development. The combination of the miRNA and their targets would certainly reveal the intricacies of regulation going on in each tissue.

In conclusion, even though ∼247 mature miRNAs have been identified for zebrafish (miRBase19) we have categorized sex specific and tissues specific candidates and this attempt paves the path for tissue biomarker discovery. Addition of 459 novel pre-miRNAs will serve as a good resource for future research to understand miRNA-mRNA regulation in zebrafish. The high degree of similarity in the protein coding genes between zebrafish and humans promotes the application of discoveries in zebrafish to humans; functional analaysis of the novel zebrafish miRNAs will have implications in understanding their role in humans.

## Methods

### RNA Isolation and Libraries Construction

Total RNA were extracted using mirVana™ miRNA Isolation Kit (Life Technologies) from independent pools of embryos, male/female adult brains, male/female adult guts, male/female adult livers, ovaries, testes, eyes and hearts (2.5, 6, 12, 24 hpf pooled RNA) and 15d juvenile fish. For each sample RNA was collected from 3 biological replicates. Concentration and quality of the RNA was analyzed on Nanodrop (A_260_/A_280_ and A_260_/A_230_ ratios) and the integrity of the RNA was verified using RNA6000 chips on a Bioanalyzer (Agilent Technologies).

Sequencing libraries for miRNA was constructed usingTruSeq® Small RNA Sample Prep Kit, Illumina. Sequencing library was prepared as per the instruction given in the manual.

### Sequencing

Sequencing was done on an Illumina HiSeq 2000 machine using TruSeq v3 cBot and SBS kits in Genome Institute of Singapore sequencing facility.

### Quantitative PCR

Quantitative PCR was performed to verify the miRNA sequencing using miRCURY LNA™ Universal RT microRNA PCR (Exiqon). 20ng of total RNA from each pool was used as starting material and qPCR was setup according to manufacturer instruction. Cycling and data acquisition were performed on an ABI, 7900HT machine. The cycling conditions consist of a 10 minutes initial denaturation at 95°C followed by 40 cycles of 95°C for 10 seconds, 60°C for 1 minute. Data was acquired at the end of each annealing/extension step. The data was analysed with comparative Ct method (2–[delta][delta]Ct) using two miRNA housekeeping genes (dre-let-7a and dre-miR-10c).

### Processing of the reads

The reads were first subjected to adapter removal through the cutadapt program (Martin 2011). The trimmed reads were then collapsed to remove redundancy and to obtain a unique sequence fasta file through the mapper module of miRDeep2 package (Friedlander et al. 2008); (Friedlander et al. 2012). The headers of the unique fasta file comprise of a running number and the frequency of the particular trimmed read (For details see **Supplemental Methods**).

### Annotation of the reads and the elimination pipeline

The unique reads fasta file was then put through an elimination pipeline module (Vaz et al. 2010) comprising of a series of sequence similarity searches with the annotated databases. At each step the reads were matched to an annotated database with a maximum of two mismatches. The matched reads were removed and the unmatched ones were further matched to the next annotated database, finally culminating into an un-annotated pool of reads that served as a source of novel miRNAs and novel sRNAs. The annotated databases used in the elimination pipeline were as follows:

i. Precursor miRNAs were obtained from miRBase, Release 19 comprising of 344 precursor miRNAs.
ii. Non coding RNA file ‘Danio_rerio.Zv9.71.ncrna.fa’ consisting of RNAs such as the miRNAs, rRNAs, sn/snoRNAs, lincRNAs and miscRNAs was downloaded from Ensembl FTP site.
iii. tRNAs (Zv9) were downloaded from http://gtrnadb.ucsc.edu/Dreri/site.
iv. The‘rna.fa’ file comprising of majorly mRNAs was downloaded from NCBI FTP site.

The reads that matched to the annotated databases were assigned to that particular class of RNA and pie-charts were constructed indicative of the distribution of the annotated sRNA classes as well as the unannotated reads in a sample.

### Known miRNA expression profile generation, normalization and clustering

The known mature miRNA expression profile was generated by using the quantifier module of the miRDeep2 package (Friedlander et al. 2012) that gives the read counts for the known miRNAs (For details see **Supplemental Methods**). The raw reads expression profile generated for all the replicates of the samples were subjected to Trimmed Mean of M-values (TMM) normalisation using the bioconductor package edgeR (Robinson et al. 2010). The normalized expression profiles for all the replicates of all the samples were then subjected to hierarchical clustering as well as PCA clustering for quality control purpose as well as to look at the similarity among the samples.

### Detection of tissue and sex specific known miRNAs

For determining the tissue specific known miRNAs, firstly, the TMM normalized mature miRNA expression profiles of all the tissues were compared to the embryo taken as control to obtain differentially expressed miRNAs (tissue vs embryo). The GLM framework in edgeR (Robinson et al. 2010) was used and the treatDGE function was applied. A log Fold Change (logFC) change cut-off of 1, representing the size of the change and a p-value with FDR cut-off of <= 0.05 representing the significance of the change was used to get the differentially expressed miRNAs.

Once the differentially expressed mature miRNAs of all the tissues (with respect to the embryo) were obtained, the upregulated mature miRNA sets were compared with each other except with its male/female counterpart to determine the miRNAs that were commonly upregulated as well as those that were only upregulated (specific) in that tissue. If a miRNA was found to be commonly upregulated, the log Fold change values of the commonly upregulated miRNA was compared by calculation of the log Fold change difference = logFC1-logFC2. The logFC1 and logFC2 are the fold change values of a commonly upregulated miRNA in the concerned tissue and the other tissue sample which it was being compared to respectively. If the log Fold change difference was less than 1, the miRNA was considered to be commonly upregulated in both the tissues. However, if the log Fold change difference was greater than 1, it was considered to be “enriched” in that concerned tissue only.

Those upregulated mature miRNAs that were “specific” to a tissue or those that were highly “enriched” in a tissue in comparison to others were called as “Tissue specific miRNAs”. For tissues that had male and female counterparts (brain, gut, liver), a union of the two sets was taken to get the complete number of tissue specific miRNAs.

For determining the sex specific known miRNAs, the TMM normalized mature miRNA expression profiles of the female counterpart of the tissue was compared to the male counterpart (female vs male) using the the GLM framework in edgeR (Robinson et al. 2010) and the treatDGE function. A log Fold Change (logFC) change cut-off of 1, representing the size of the change and a p-value with FDR cut-off of <= 0.05 representing the significance of the change was used to get the differentially expressed miRNAs. These differentially expressed miRNAs among the male and female were called as “Sex specific miRNAs”.

### Novel miRNA prediction pipeline

The un-annotated sequences left after the elimination pipeline, were used for the novel miRNA prediction. For this purpose, the un-annotated sequence files of the replicates of a particular tissue were combined and first mapped to the zebrafish genome using mapper.pl module and then subjected to the miRDeep2.pl module (Friedlander et al. 2012) to obtain the novel miRNAs (For details see **Supplemental Methods**).

### Detection of tissue specific novel pre-miRNAs

The novel pre-miRNA sequences of all the tissues predicted by miRDeep2 were compared with each other including the embryo sample using an in-house sequence similarity search perl program that looks for equal sequence matches as well as substrings and overlaps to obtain the total matched set of novel pre-miRNA sequences. The total matched set was then subtracted from the total predicted set of novel pre-miRNA sequences to obtain the unmatched set of novel pre-miRNA sequences. The unmatched set of novel pre-miRNA sequences were deemed as tissue specific novel pre-miRNAs for each tissue and embryo specific novel pre-miRNAs for the embryo **(Supplemental Figure S4)**.

The entire protocol comprising of processing of the reads, quantification of known miRNAs, elimination pipeline and novel miRNA prediction has been depicted in **Figure 1.**

## Data Access

The data discussed in this publication have been deposited in NCBI’s Gene Expression Omnibus and are accessible through GEO Series accession number GSE57169. (http://www.ncbi.nlm.nih.gov/geo/query/acc.cgi?acc=GSE57169).

## Acknowledgements

Financial support received from Molecular Genomics (Singapore), LKC School of Medicine, NTU, (Singapore) and ASTAR (Singapore) are gratefully acknowledged.

## Disclosure Declaration

The authors declare that they have no competing interests.

## Supplemental Figures

**Supplemental Figure S1 – Principal Component Analysis plot depicting the clustering of the samples.**

The three biological replicates of all the tissue samples including embryo clustered closely indicating less experimental variation among the replicates. The tissue samples having male and female counterparts like the brain, gut and liver showed proximity among their male and female counterparts, indicating that the variations caused due to sex was less compared to the variations caused by the tissue.

**Supplemental Figure S2 – MiRDeep2 displays a high sensitivity for identification of known miRNAs from zebrafish.**

The number of known miRNAs picked up by miRDeep2 in comparison to the total number of miRNAs in the sample; at the similar cut-off used for novel miRNA prediction was used as an indicator of its sensitivity. The sensitivity of miRDeep2 ranged from 89 to 95% with the exception of one female liver sample, for which the sensitivity was 81%.

**Supplemental Figure S3 – Structure of a predicted novel miRNA from miRdeep2 with its aligned read sequences.**

MiRdeep2 gives as output the pdfs of the structure of the predicted novel miRNA along with the reads mapping to its mature, star and loop sequences. The top left corner has the score distribution for the predicted novel miRNA, the top right corner has the predicted hairpin structure for the novel pre-miRNA and the major part comprises of the alignment of the reads to the mature (red), loop (yellow) and the star regions (purple) of the precursor miRNA. For each read mapped, its frequency value, the number of mismatches with which it maps, and the sample it belongs to is given at the bottom right.

**Supplemental Figure S4 – Schematic representation of obtaining specific novel pre-miRNAs for embryo.**

The coloured circles represent the set of predicted novel pre-miRNAs for each tissue sample. The 78 predicted novel pre-miRNAs of embryo were compared with the set of predicted novel pre-miRNAs of other tissues to find the ones that matched. The total set of matched novel pre-miRNAs were 59. Therefore the unmatched set of 19 was considered as specific novel pre-miRNAs for embryo. This procedure was followed for the other tissue samples to obtain the novel pre-miRNAs specific to them.

### Supplemental Tables

**Supplemental Table S1 – Differentially expressed known miRNAs in the tissue samples with respect to embryo.**

To determine the tissue specific known miRNAs, the TMM normalized expression profiles of the known mature miRNAs of all the tissues were first compared with those of the embryo samples; using GLM edgeR analysis to obtain the differentially expressed miRNAs (DE miRNAs). A log Fold Change (logFC) change cut-off of 1, representing the size of the change and a p-value with FDR cut-off of <= 0.05 representing the significance of the change was used to get the differentially expressed known mature miRNAs.

**Supplemental Table S2 – List of the tissue specific known miRNAs.**

The up-regulated DE known mature miRNA sets of all the tissues were compared with each other except with its male/female counterpart to obtain the up-regulated DE mature miRNAs only found or the most enriched in that particular tissue to obtain tissue specific miRNAs. The union of the tissue specific miRNAs for the male and female counterparts of brain, gut and liver were taken as their respective tissue specific miRNAs. The brain had the highest number of known tissue specific miRNAs (36).

To obtain embryo specific known miRNAs, the down-regulated DE known mature miRNA sets of all the tissues were compared to obtain the commonly down regulated miRNAs. These commonly down regulated miRNAs were called “embryo specific” (9).

**Supplemental Table S3 – List of the sex specific known miRNAs.**

To determine the sex specific known miRNAs, the TMM normalized mature miRNA expression profiles of known miRNAs of the male and female counterparts of the brain, gut and liver were compared using the GLM edgeR analysis to obtain differentially expressed miRNAs (DE miRNAs). A log Fold Change (logFC) change cut-off of 1, representing the size of the change and a p-value with FDR cut-off of <= 0.05 representing the significance of the change was used to get the differentially expressed miRNAs. The DE mature miRNAs among the male and female counterparts were deemed as the sex specific miRNAs. The liver had the highest number of sex specific miRNAs (34).

**Supplemental Files**

**Supplemental File S1 –** Raw and TMM Normalized known mature miRNA expression profiles of all the samples. (Excel file with 2 sheets. Sheet 1: Raw, Sheet 2: TMM Normalized)

**Supplemental File S2 –** Details of the differentially expressed (DE) known mature miRNAs of all the tissue samples with respect to embryo including the logFC, logCPM, P-value and FDR values. (Excel file with 5 sheets. Sheet1: male-female brain DE miRNAs, Sheet2: male-female gut DE miRNAs, Sheet3: male-female liver DE miRNAs, Sheet4: ovary-testis DE miRNAs, Sheet5: eye-heart DE miRNAs).

**Supplemental File S3 –** Details of the tissue specific known miRNAs of embryo and all the tissues including the logFC values. (Excel file with 8 sheets. Sheet1: embryo specific known miRNAs, Sheet2: brain specific known miRNAs, Sheet3: gut specific known miRNAs, Sheet4: liver specific known miRNAs, Sheet5: ovary specific known miRNAs, Sheet6: testis specific known miRNAs, Sheet7: eye specific known miRNAs, Sheet8: heart specific known miRNAs).

**Supplemental File S4 –** Details of the sex specific known miRNAs of brain (Sheet1), gut (Sheet2), liver (Sheet3) (female vs male counterparts) and differentially expressed miRNAs of ovary vs testis (Sheet4), including the logFC, logCPM, P-value and FDR values. (Excel file with 4 sheets).

**Supplemental File S5 –** Details of the predicted novel miRNAs of all the samples including the IDs, miRDeep2 scores, read counts, precursor and mature sequences and precursor coordinates. (Excel file with 11 sheets. Sheet1: embryo predicted novel miRNAs, Sheet2: male brain predicted novel miRNAs, Sheet3: female brain predicted novel miRNAs, Sheet4: male gut predicted novel miRNAs, Sheet5: female gut predicted novel miRNAs, Sheet6: male liver predicted novel miRNAs, Sheet7: female liver predicted novel miRNAs, Sheet8: ovary predicted novel miRNAs, Sheet9: testis predicted novel miRNAs, Sheet10: eye predicted novel miRNAs, Sheet11: heart predicted novel miRNAs.

**Supplemental File S6 –** Precursor sequences of the 459 predicted novel miRNAs along with their 5p and 3p mature miRNAs. The novel predicted miRNAs are given the prefix : “gis-dre-mir” (Text file).

**Supplemental File S7 –** Details of the tissue specific predicted novel pre-miRNAs including the IDs, miRDeep2 scores, read counts, precursor and mature sequences and precursor coordinates. (Excel file with 9 sheets. Sheet1: embryo specific novel miRNAs, Sheet2: brain specific novel miRNAs, Sheet3: gut specific novel miRNAs, Sheet4: liver specific novel miRNAs, Sheet5: ovary specific novel miRNAs, Sheet6: testis specific novel miRNAs, Sheet7: eye specific novel miRNAs, Sheet8: heart specific novel miRNAs, Sheet9: summary).

## Authors Contributions

SM conceived the project, CWW & GPSL performed the experiments, CV and VT analyzed the data and SM, PWI, VT & CV wrote the paper.

